# Resisting the resistance: The antimicrobial peptide DGL13K selects for small colony variants of *Staphylococcus aureus* that show increased resistance to its stereoisomer LGL13K, but not to DGL13K

**DOI:** 10.1101/2024.02.26.582211

**Authors:** Sven-Ulrik Gorr

## Abstract

About 30% of the population are nasal carriers of *Staphylococcus aureus*, a leading cause of bacteremia, endocarditis, osteomyelitis, skin and soft tissue infections. Antibiotic-resistant bacteria, in particular, are an increasing problem in both hospital and community settings. In this study, we sought to determine the cellular consequences of long-term exposure of *S. aureus* to the antimicrobial peptide stereo-isomers, DGL13K and LGL13K. Both peptides selected for mutations in the chorismate/menaquinone biosynthetic pathway, which resulted in increased resistance to LGL13K but not DGL13K. DGL13K-selected isolates showed a mutation in *aroF*, while *menA* and *menH* were mutated in LGL13K-selected isolates. The latter also contained a mutation of *frsA*. The peptide-selected isolates exhibited golden coloration, suggesting increased production of the carotenoid staphyloxanthin, which could enhance resistance to cationic AMPs. The peptide-selected isolates grew as small colony variants, which have also been associated with resistance to AMPs. Addition of menaquinone to the growth medium reduced the generation time of DGL13K-selected mutants, but not LGL13K-selected mutants. Instead, the latter showed greatly reduced ATP-levels, suggesting defective electron transport, which is also associated with menadione auxotrophism. The mechanisms behind the differential effect of DGL13K and LGL13K on *S. aureus* resistance remain to be elucidated. The finding that DGL13K induced resistance to the stereo-isomer LGL13K but not to DGL13K itself, suggests that peptide primary structure is responsible for inducing bacterial defense mechanisms but the peptide secondary structure determines if the defense mechanisms are effective against each peptide.

## Introduction

Mucosal surfaces are exposed to both commensal and invading bacteria, which must survive in an environment rich in host-defense molecules, including antimicrobial peptides (AMPs) (1). As an example, about 30% of the population are nasal carriers of *Staphylococcus aureus*, which causes no harm in most people (2). Thus, in one study, only about 1% of nasal carriers developed bacteremia but over 80% of *S. aureus* isolated in bacteremia were of endogenous nasal origin (3). A highly adaptable metabolism allows *S. aureus* to infect various host environments (4) and it is a leading cause of bacteremia, endocarditis, osteomyelitis, skin and soft tissue infections. Antibiotic-resistant bacteria, in particular, are an increasing problem in both hospital and community settings. As an example, the U.S. Centers for Disease Control and Prevention estimate that over 300,000 infections and 10,000 deaths annually are attributable to methicillin-resistant *S. aureus* (MRSA), which is listed as a serious threat by the agency (5). WHO also lists *S. aureus* as a high priority pathogen for which novel antibiotics are urgently needed (6)

A dozen cationic AMPs mediate the antibacterial function of human nasal fluid (7) and 45 AMPs have been identified in the oral cavity (8). Bacteria use a variety of mechanisms to defend against AMPs, including electrostatic repulsion, cell wall alteration, membrane alteration, proteolysis, protein binding and efflux pumps (9-12). Accordingly, a number of resistance genes have been identified in Gram positive bacteria (13), including two dozen resistance genes - the “resistome”, which are associated with resistance to the AMP LL-37 (14). The identification of diverse resistance genes, belonging to different functional families, points to the plasticity of bacterial resistance. As an example, the Gram-positive bacteria *Enterococcus faecalis* and *Streptococcus gordonii* are resistant to the AMP LGL13K. However, deletion of *dltA*, which functions in D-alanylation of teichoic acids, renders the bacteria sensitive to LGL13K. Upon prolonged exposure to LGL13K, these *dltA*^*-*^ bacteria develop de novo resistance to this peptide through a separate mechanism (15).

Host-defense peptides have served as inspiration for the design of therapeutic AMPs (9, 16). To be successful, a therapeutic AMP must overcome multiple resistance mechanisms, which has raised the concern that resistance to therapeutic AMPs could also render bacteria resistant to host-defense peptides (“arming the enemy”) (17, 18). Indeed, cross-resistance has been achieved experimentally in vitro and in animal models (19-21) but this is not a consistent outcome of AMP selection, which may increase, decrease or have no effect on the minimum inhibitory concentration (MIC) of a different antimicrobial peptide; reviewed in (9). Moreover, the wide-spread use of nisin and polymyxin B, without generalized resistance, suggests that this is not a general threat to host-defenses (22).

We designed the AMP LGL13K, which kills Gram-negative bacteria and their biofilms (23, 24). However, LGL13K is not effective against Gram-positive bacteria and is susceptible to proteolysis. To overcome this problem, an all-D-amino acid isomer, DGL13K, which resists proteolysis, was designed (23). DGL13K is highly effective against Gram-positive bacteria and this activity is independent of proteolytic activity (15). In this report, we extend these studies to *S. aureus* and show that these Gram-positive bacteria are also resistant to LGL13K but not DGL13K. Remarkably, either peptide isomer increases resistance to LGL13K but not DGL13K. The resistant bacteria contain mutations in the shikimate/menaquinone biosynthetic pathways and exhibit a small colony variant (SCV) phenotype that differs from traditional SCVs.

## Materials and Methods

### Bacterial cultures

*Pseudomonas aeruginosa* Xen41, a bioluminescent derivate of *P. aeruginosa* PAO1, was obtained from Xenogen (Alameda, CA; now Perkin-Elmer, Waltham., MA). *Staphylococcus aureus* Xen36, a bioluminescent derivate of America Type Culture Collection strain 49525, was obtained from Perkin-Elmer. *P. aeruginosa* and *S. aureus* were cultured from glycerol stocks at 37°C with constant shaking at 200 rpm in liquid cultures prepared in Luria-Bertani broth and Todd Hewitt Broth (THB) (Difco, Franklin Lakes, NJ), respectively. Overnight cultures typically reached an optical density at 600 nm (OD600) corresponding to 5 x 10^8^ CFU/ml for *P. aeruginosa* and 3 x 10^8^ CFU/ml for *S. aureus*. Colonies of the latter were isolated on 1.5% agar prepared in THB and cultured at 37°C.

### Peptides

LGL13K and DGL13K were purchased from AappTec (Louisville, KY) or Bachem (Torrance, CA) at >95% purity. Peptide identity and purity were confirmed by the supplier by mass spectrometry and RP-HPLC, respectively. Peptides were dissolved at 10 mg/ml in sterile 0.01% acetic acid and these stock solutions stored at 4°C until use. Peptide batches were tested for antimicrobial activity by MIC assays. The stock solutions retained activity for at least 2 years at 4°C (**Fig. 1**).

**Figure 1.**
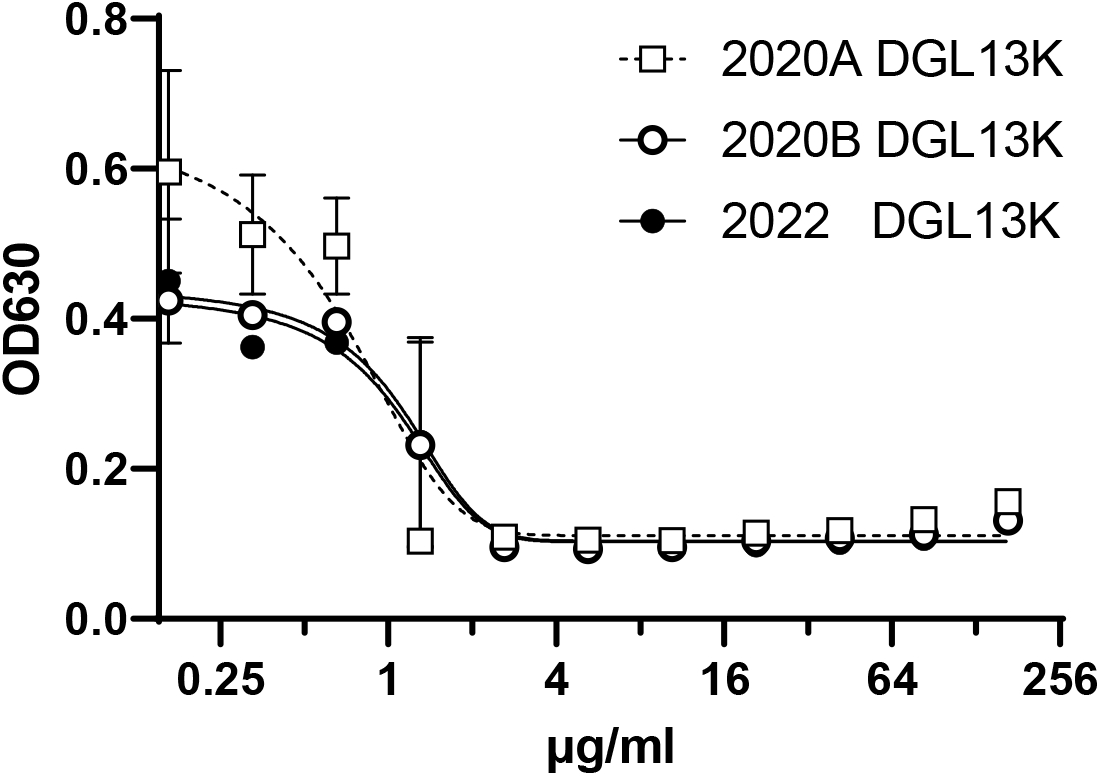
DGL13K retains activity in solution. Two stock solutions of DGL13K were stored at 4°C for 2 years (2020A and B, □ o) or freshly made (2022 ●). MIC was tested against S. aureus Xen 36. Data are shown as mean ± range of duplicate samples, N=2.

### Minimum Inhibitory Concentration

MIC assays were performed essentially as previously described (15, 25). The overnight cultures were diluted to about 10^5^ CFU/ml in THB for *S. aureus* and Mueller-Hinton Broth (Difco) for *P. aeruginosa*. Diluted bacteria (100 µl/well) were incubated in 96-well polypropylene microtiter plates with two-fold serial dilutions of each peptide in 20 µl ‘10% PBS’ (a 10-fold dilution of PBS in sterile water). LGL13K was tested in the concentration range 1667 µg/ml to 1.6 µg/ml while DGL13K was tested in the range 167 µg/ml to 0.16 µg/ml. Control samples without added peptide were included in each assay. Culture plates were incubated overnight at 37°C with constant shaking on a Stovall Belly dancer lab shaker at speed 5 (IBI, Dubuque, IA). The OD was determined at 630 nm (OD630) in a BioTek Synergy HT plate reader (BioTek, Winooski, VT; now Agilent, Santa Clara, CA). The OD630 for each peptide dilution was plotted and the MIC determined in four parallel replicates.

For serial MIC assays, a single representative well, containing a peptide concentration 2-fold lower than the MIC (0.5xMIC), was sampled and the bacteria diluted 1100x in THB or Mueller-Hinton Broth and used to inoculate the next MIC assay. Sampling in two separate experiments was repeated for 6 days for *S. aureus* and 12 days for *P. aeruginosa*.

### Peptide-selected isolates

Aliquots of *S. aureus* cultures, that were treated with DGL13K or LGL13K in 5-6 consecutive MIC assays, were streaked on THB agar or cultured overnight in the absence of DGL13K or LGL13K and then streaked on THB agar. Individual colonies were selected and expanded in THB, in the absence of DGL13K or LGL13K, overnight at 37°C with shaking at 200 rpm. The overnight cultures of individual colonies were mixed with glycerol (10% final concentration) and stored at -80°C as *DGL13K-selected and LGL13K-selected isolates*, respectively.

### Genome sequencing

Bacterial pellets of peptide-selected isolates were submitted to the University of Minnesota Genomics Center for DNA purification, genomic library generation and MiSeq genomic sequencing. gDNA samples were converted to Illumina sequencing libraries using Illumina’s DNA Prep Sample Preparation Kit (Illumina, San Diego, CA). Briefly, 1-500 ng of gDNA was simultaneously fragmented and tagged with a unique adapter sequence. The DNA was simultaneously indexed and amplified by PCR. Final library size distribution was validated using capillary electrophoresis and quantified using PicoGreen fluorimetry.

Libraries were sequenced on an Illumina MiSeq platform (2x300 bp) using Illumina’s SBS chemistry. Primary analysis and de-multiplexing were performed using Illumina’s bcl-convert v4.0.3. The resulting FASTQ files were mapped to the published *S. aureus* reference genome sequence (NCBI Reference Sequence: NC_002950.2) via BWA (0.7.17-r1188) to generate BAM files. Variant calling was done in parallel across all samples via Freebayes using a minimum variant frequency of 0.01 and minimum coverage of 34 reads. Polymorphism frequencies in each culture were determined and gated at >10% threshold. Raw data files associated with genome sequencing are maintained by the Minnesota Supercomputing Institute (https://www.msi.umn.edu/).

### Growth curves and colony size

Aliquots (5 µl) of glycerol stocks of peptide-selected isolates were diluted in 1 ml THB and 200 µl aliquots cultured in 96-well plates at 37°C without shaking. OD_630_ was read at 45-90 min intervals and fitted to an exponential growth curve, which was used to calculate generation time (doubling time) (Graphpad Prism 8; GraphPad Software. Boston, MA).

To visualize differences in colony size, overnight cultures of wild-type and peptide-selected isolates were diluted in THB and an aliquot spread on THB agar and again cultured overnight at 37°C.

### Colony pigmentation

To determine colony pigmentations, aliquots of wild-type or peptide–selected isolates were plated on blood agar containing 50 µg/l menadione, 5 mg/l hemin, and 0.25% sheep blood. The plates were incubated 2 days at 37°C.

### Menaquinone supplementation

To determine if menaquinone supplementation affected growth rate, wild-type and peptide-selected isolates were cultured in Mueller-Hinton broth alone or supplemented with 50 µM menaquinone (Vitamin K2; Supelco), which was added from a 5mM stock solution in 95% ethanol. Exponential growth curves were determined and the generation time calculated, as described above.

### Cellular ATP content

*S. aureus* Xen 36 contain a stable copy of the modified *Photorhabdus luminescens* luxABCDE operon, which utilizes cellular ATP to emit light. We employed this system to determine cellular ATP content, as a measure of metabolic activity (26). Bacteria were cultured to stationary phase and the bacterial luminescence (Relative Light Units - RLU) was recorded and expressed relative to OD600 of the culture, to account for different growth rates.

## Results

### Resistance to LGL13K or DGL13K

The Gram-negative bacteria *P. aeruginosa* are highly susceptible to both LGL13K and DGL13K, showing only a 2-4-fold difference in MIC between the two peptide isomers (24, 27, 28). In fact, in a serial peptide exposure experiment, the MIC only increased about 2-fold for LGL13K and this increase was not statistically significant, suggesting that the bacteria are not able to mount resistance to either peptide stereoisomer (27). To validate this finding, *P. aeruginosa* was serially exposed to LGL13K or DGL13K. As expected, neither LGL13K nor DGL13K showed an increased MIC after serial exposure for up to 12 days (**Fig. 2A**).

**Figure 2.**
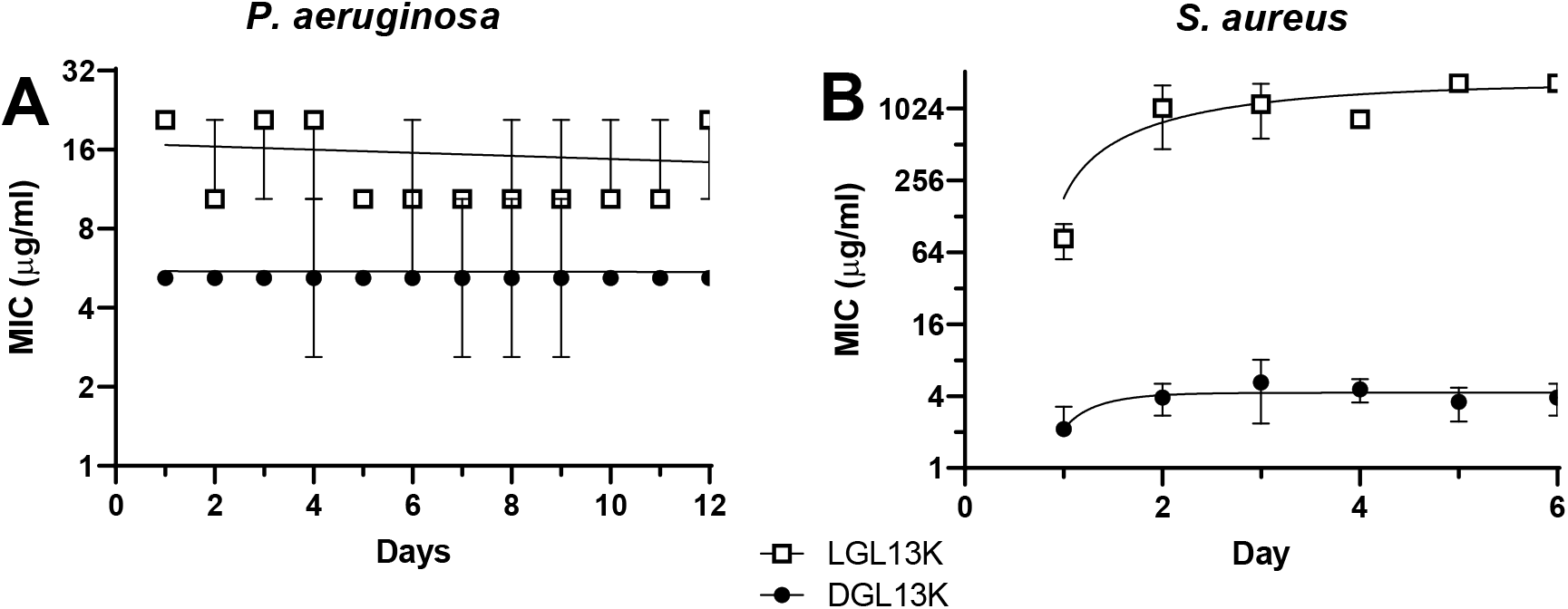
Acquired resistance of P. aeruginosa (A) and S. aureus (B) after serial MIC assays for 12 and 6 days, respectively. Bacteria were treated with LGL13K (□) or DGL13K (●) in successive MIC assays, which were each inoculated with the culture at 0.5xMIC from the previous day. Daily MICs are shown as the mean ± 95%CI of two independent experiments, each performed in quadruplicate (N=8).

In contrast to *P. aeruginosa*, Gram-positive bacteria are relatively resistant to LGL13K, but not DGL13K (15, 28). To characterize further this stereo-selective resistance, these studies were extended to the Gram-positive bacteria *S. aureus*. This species shows a lower initial MIC to LGL13K than the previously tested Gram-positive species (15, 28), which allows for further selection with LGL13K. **Figure 2B** shows that the MIC for LGL13K rapidly increased during selection, reaching a 20-fold increase on day 2-3. In contrast, the initial MIC for DGL13K was about 40-fold lower than that for LGL13K and increased less than 2-fold over six days of selection (**Fig. 2B**), consistent with our results with other Gram-positive bacteria (15).

Bacteria selected with either DGL13K or LGL13K for 5-6 days were cultured in the absence of peptide and individual colonies isolated for further study (peptide-selected isolates)

### Cross-over resistance

To further explore the different levels of resistance induced by the two stereo-isomers, the peptide-selected isolates were tested against both stereo-isomers (**Fig. 3**). The DGL13K selected isolates remained susceptible to DGL13K, consistent with the selection experiments shown in Fig. 2B. Similarly, the LGL13K-selected isolates retained an increased MIC for this peptide, compared to WT bacteria (**Fig. 3**). In contrast, the LGL13K-selected isolates were readily killed by DGL13K, suggesting that resistance was selective for the L-isomer of this peptide. Surprisingly, the DGL13K-selected isolates became resistant to LGL13K, although they had not previously been exposed to this stereo-isomer (**Fig. 3**). These results suggest that both peptide stereo-isomers induce similar resistance mechanisms but these mechanisms are not effective against the D-isomer, which “resists the resistance”.

**Fig. 3.**
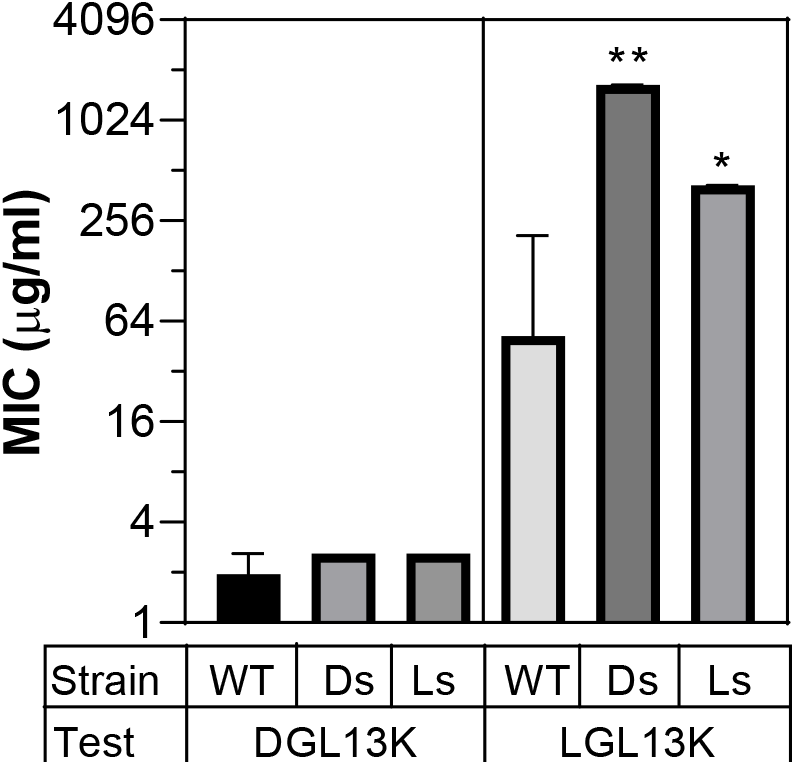
Cross-over MIC assays. Wild-type (WT), DGL13K-selected (Ds) or LGL13K-selected (Ls) isolates were analyzed in MIC assays against the test peptide, DGL13K or LGL13K. The MIC was determined in 2-5 independent experiments and expressed as median MIC ± 95%CI. ^*^) different from WT, P<0.002; ^**^) different from WT, P<0.0001; N=6-21.

### Genome sequencing

To investigate the molecular mechanism behind the increased MIC for LGL13K, DGL13K-selected and LGL13K-selected isolates of *S. aureus* were submitted for whole genome sequencing. The analysis was focused on mutations that distinguished the peptide-selected isolates from the *S. aureus* Xen 36 wild-type genome. As shown in **Table 1**, four different mutations were identified in the isolates. Three of the four mutations mapped to the biosynthetic pathways for chorismate and menaquinone (KEGG: map00400 and map00130). DGL13K-selected isolates were mutated in *aroF* (DAHP synthase), which catalyzes the first step in the synthesis of chorismate (shikimate pathway) (29). LGL13K-selected isolates were mutated in *menA* or *menH*, which act in the conversion of chorismate to menaquinone. *menH* mutants also showed mutation of *frsA*, which acts as a switch between respiration and fermentation (30). Notably, these mutations appear to be novel and were not identified in a recent screen for the AMP-induced resistome in *S. aureus* (14).

**Table 1.** Genome sequencing of peptide selected isolates. Bacteria were selected with LGL13K (two selection experiments) or DGL13K (one selection experiment) and four individual colonies from each set were sequenced. The mutated genes, nucleotide (NT) change and the corresponding change of protein sequence (AA) are shown.

| Selection | GENE | NT change | AA change |
| --- | --- | --- | --- |
| **LGL13K1** | *frsA* | A->T | Tyr256Asn |
|  | *menH* | A->T | Ile106Leu |
| **LGL13K2** | *menA* | C->G | Ala91Pro |
| **DGL13K1** | *aroF* | G->A | Arg210Cys |

### Characterization of small colony variants

The peptide-selected isolates showed colony morphology reminiscent of SCVs (**Fig. 4A**), which are associated with antibiotic resistance (31-34). The LGL13K-selected colonies were somewhat smaller than WT colonies, while DGL13K-selected colonies were substantially smaller than WT colonies. This difference was quantified from exponential growth curves. The generation time (doubling time) for DGL13K-selected isolates was 3.5-fold longer than for wild-type cultures, while LGL13K-selected isolates showed a 1.8-fold increase in doubling time (**Fig. 4B**), consistent with the observed colony morphology.

**Figure 4.**
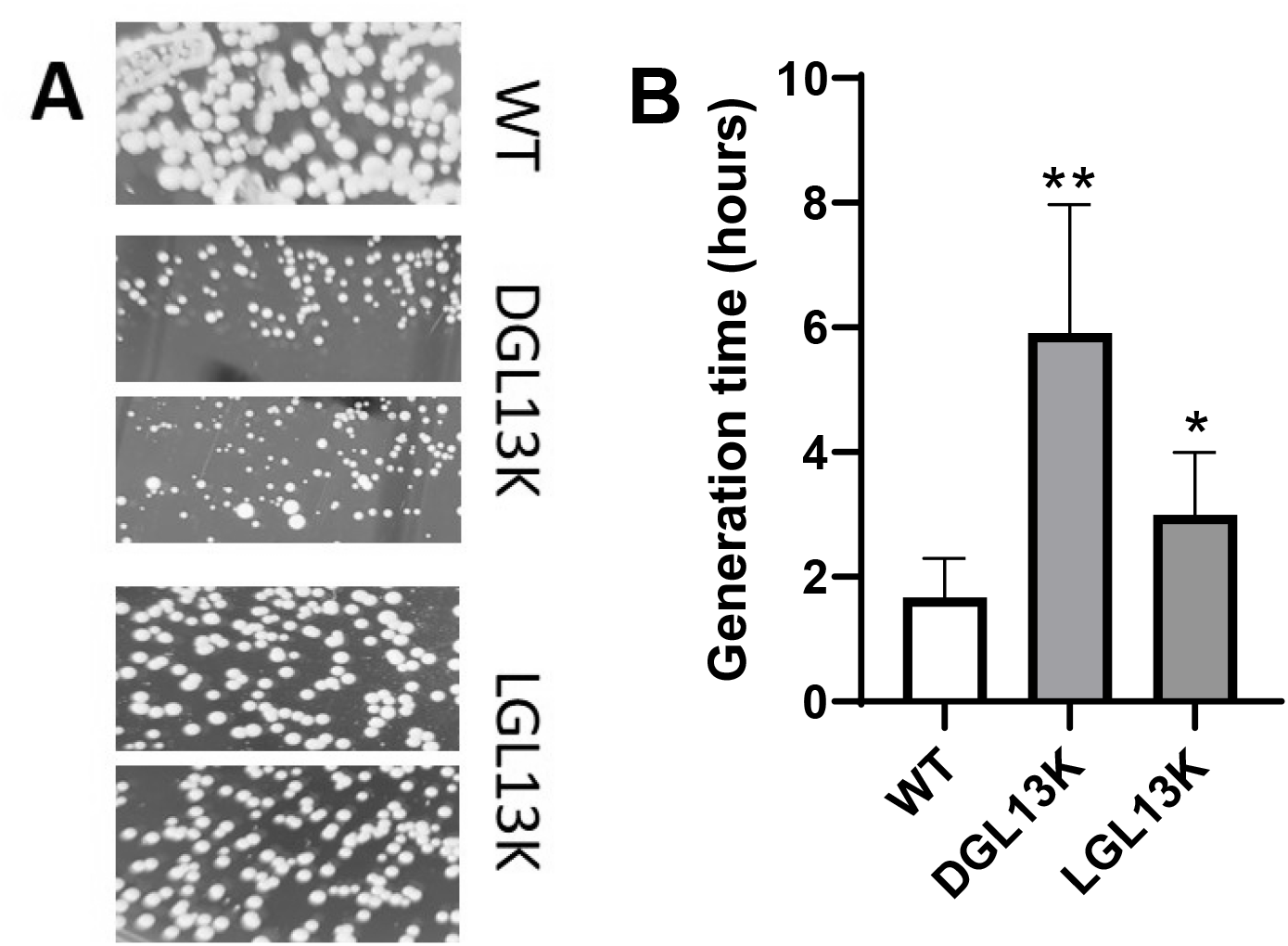
Growth characteristics of peptide-selected isolates. **A**. Wild-type (WT), and two isolates each selected with DGL13K or LGL13K were plated on THB agar and photographed to show the relative colony sizes. The images are representative of three independent cultures. **B**. The generation time (doubling time) was calculated from exponential growth curves for WT, DGL13K-selected and LGL13K-selected isolates. Data from four independent experiments with 1-2 WT samples and 3-6 isolates are shown as mean ± 95%CI, N = 6-21. ^*^) different from WT, P<0.05; ^**^) different from WT, P<0.001.

In addition to longer generation times and the resulting small colony size, small colony variants of *S. aureus* have been characterized by a lack of pigmentation and auxotrophism caused by mutations in metabolic pathways that lead to defective electron transport and decreased ATP production (32). *S. aureus* Xen36 is a derivative of ATCC 49525 and produces non-pigmented (cream colored) colonies (35). The lack of pigmentation of WT Xen36 was confirmed by culture on blood agar for 2 days (**Fig. 5A**). In contrast, the DGL13K-selected variants showed strong pigmentation while LGL13K-selected variants showed varying levels of pigmentation (**Fig. 5A**). Thus, higher levels of pigmentation were correlated with smaller colony size.

**Figure 5.**
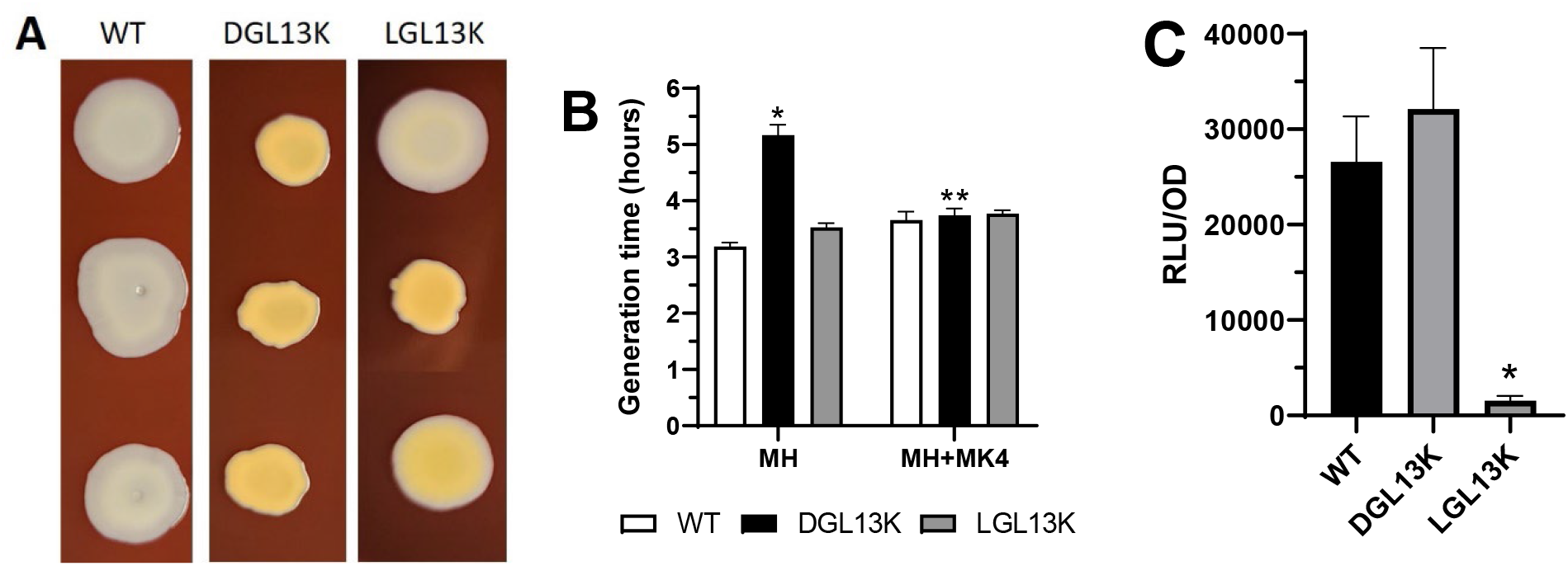
Characterization of peptide-selected isolates. **A**. Aliquots of WT, DGL13K-selected and LGL13K-selected isolates were cultured on blood agar for two days. **B**. Generation time (doubling time) for WT (white bar), DGL13K-selected (black bar) and LGL13K selected (grey bar) isolates cultured in Mueller-Hinton broth with (MH+MK4) or without (MH) menaquinone supplementation. Data from two independent experiments are show as mean ± 95%CI, ^*^) different from MH WT and LGL13K, ^**^) different from MH DGL13K. P<0.0001, N=3-7. **C**. ATP activity in peptide-selected isolates. Bacterial luminescence (RLU) was recorded at stationary phase growth and expressed relative to optical density of the culture at 600 nm (OD). The average OD in each group was WT=0.7; DGL13K=0.5; LGL13K=0.6. The data from eight independent experiments are shown as mean ± 95%CI. ^*^) different from WT and DGL13K, P<0.0001, N=26-35.

SCVs are typically auxotrophic including menadione auxotrophy (36, 37), which makes the bacteria unable to synthesize menaquinone (38). Given the mutations identified in the chorismate and menaquinone biosynthetic pathways (Table 1), we tested the growth rates of peptide-selected isolates cultured in the presence or absence of menaquinone. The generation times of WT and LGL13K-selected isolates were not affected by this supplementation, while the longer generation time of DGL13K-selected isolates was reduced to that of the other samples in the presence of menaquinone (**Fig. 5B**).

Menadione-auxotrophy is associated with electron transport deficiency that reduces ATP production (37). *S. aureus* Xen36 have been engineered to express a *Photorabdus luminescens* luciferase, which can be used as an indicator of cellular ATP levels. To determine if peptide-selected isolates showed defects in ATP production, the bacteria were cultured to stationary phase and the relative ATP-dependent luminescence was recorded. **Fig. 5C** shows that the DGL13K-selected isolates expressed similar ATP levels as WT bacteria while the LGL13K-selected isolates were highly deficient in ATP content.

## Discussion

We have previously reported that *S. aureus* are relatively resistant to the AMP LGL13K but not the stereo-isomer DGL13K (28). In this study, we sought to determine the cellular consequences of this difference. Surprisingly, both peptides selected for mutations in the chorismate/menaquinone biosynthetic pathway, which resulted in increased resistance to LGL13K but not DGL13K. Thus, bacteria selected with DGL13K become resistant to the stereo-isomer LGL13K but DGL13K can evade this resistance (“*resisting the resistance*”). Cross-resistance between different AMPs has been observed experimentally (19-21). Indeed, selection of bacteria resistant to one AMP may increase, decrease or have no effect on the MIC of a different AMP; reviewed in (9). This has led to the proposal that wide-spread use of AMPs could render the host-defense peptides ineffective against invading bacteria by “arming the enemy” (17, 18). However, it has been countered that the AMPs nisin and polymyxin have been in general use for decades without affecting host-defenses (22). The present results suggest that resistance to an AMP can be overcome by a closely related AMP, which bodes well for future clinical use.

Unlike early suggestions that AMPs, that act on the cell membrane, are unlikely to cause extensive resistance (39, 40), the present results clearly support subsequent findings that membrane active peptides, including LGL13K (41), can cause resistance in *S. aureus* (42-44). Our genome sequencing data suggest that the chorismate/menaquinone pathway plays a role in resistance to LGL13K. Indeed, this pathway has previously been associated with activity and resistance to AMPs (45, 46) and *menF* is part of a mutated gene complex in pexiganan-selected *S. aureus* (21). On the other hand, *men* genes were not included in the “resistome” identified upon selection of *S. aureus* with the endogenous AMP LL-37 or engineered LL-37 derivatives (14).

Both peptide isomers selected for mutations in the chorismate/menaquinone pathway. The DGL13K-selected isolates showed a mutation in *aroF* (DAHP synthase), which catalyzes the first step in the synthesis of chorismate (shikimate pathway) (29, 47). Based on the structures of DAHP synthase in other species, the conserved Arg residue at position 210 contributes to the substrate-binding site for phosphoenolpyruvate (48). It remains to be determined if this mutation affects substrate binding.

Menaquinone is synthesized from chorismate in the vitamin K pathway, which includes the genes *menA* and *menH* that were mutated in LGL13K-selected isolates. The mutation identified in *menH*, Ile106Leu, is conservative and not located in the catalytic triad or a conserved region of the protein (49). Thus, the functional significance of this mutation is unclear. However, these mutants also showed mutation of *frsA*, which acts as a switch between respiration and fermentation (30) and could affect ATP production. Indeed, ATP content is greatly reduced in the LGL13K-selected isolates (**Fig. 5C**).

In addition to mutations in the menaquinone pathway, the peptide-selected isolates exhibited golden coloration, a sign of increased production of the carotenoid staphyloxanthin, which has been assigned a protective effect against oxidative stress (50). The golden colony color was most pronounced in the DGL13K-selected isolates, suggesting that the *aroF* mutation, in particular, affected staphyloxanthin production. Increased staphyloxanthin production is associated with increased membrane rigidity and it has been suggested that this could enhance resistance to cationic AMPs (51). Indeed, the golden isolates showed increased resistance to LGL13K while DGL13K was able to overcome this barrier, suggesting that the structure of the D-stereoisomer allows it to by-pass the more rigid membrane structure. It has previously been reported that LGL13K and DGL13K differ in their interaction with components of the Gram-positive cell wall (15, 52). The present results suggest that this concept extends to interaction with components of the bacterial cell membrane. Experiments with model membranes have found that LGL13K forms a beta-sheet structure in the presence of negatively charged liposomes and destabilizes the membrane by removing lipid micelles (53). Harmouche et al. (2017) similarly found that LGL13K transitions from random coil in solution to beta-sheet conformation in the presence of negatively charged lipid membrane, which leads to deformation of the membrane and opening of the bilayer (41). The ability to form beta-sheets and self-assembled nanostructures in solution is faster for DGL13K than LGL13K and is lacking in a randomized GL13K sequence. Thus, these nanostructures correlate with the antimicrobial activity of each peptide (54). Together, these results support the notion that L-and D-isomers of AMPs are not perfect mirror images but display subtle differences in secondary structure that manifest in differentiated antimicrobial activity, beyond their differences in proteolytic susceptibility (15, 55).

The peptide-selected isolates displayed as SCVs, which have been associated with antimicrobial resistance (31, 33), including resistance to AMPs (34, 56). This colony morphology was most pronounced for the DGL13K-selected *aroF* mutants, but still notable for the LGL13K-selected mutants. In addition to colony morphology, SCVs are frequently characterized by auxotrophism for menadione, hemin and/or thymidine (37). Mutations in *menC, menD, menE*, or *menF* block the biosynthesis of menadione, which renders the bacteria unable to synthesize menaquinone (38). Indeed, addition of menaquinone to the growth medium, reduced the generation time of DGL13K-selected mutants, but not LGL13K-selected mutants, to match the faster growth rate of wild-type cells. Conversely, LGL13K-selected mutants, but not DGL13K-selected mutants, showed greatly reduced ATP-levels, suggesting that the former exhibit defective electron transport, which is associated with menadione auxotrophism (37).

The SCVs identified in this study differ from “conventional” SCVs in several ways (**Table 2**). SCVs typically present as small colonies with defects in electron transport that leads to reduce electrochemical gradient (resistance to cationic antibiotics, including AMPs), decreased ATP levels and cellular growth, and decreased pigment formation (37). The DGL13K-selected isolates present with a small colony morphology and resistance to LGL13K but not DGL13K. The ATP content is not affected but the colonies show increased pigment formation. LGL13K-selected isolates show moderately reduced colony size and greatly reduced ATP content. However, the resulting colonies only display a moderate increase in pigment formation.

**Table 2.** Properties of small colony variants selected with DGL13K or LGL13K.

|  | **Conventional SCV** | **DGL13K-selected** | **LGL13K-selected** |
| --- | --- | --- | --- |
| Colony size | small | small | intermediate |
| Growth rate | Slow | Slow (6 x WT) | Decreased (2 x WT) |
| Resistance | Cationic AMP resistance | LGL13K, increased<br>DGL13K, no effect | LGL13K, increased<br>DGL13K-no effect |
| Mutated genes |  | *aroF* | *menA, menH, frsA* |
| Pathway | N/A | shikimate | menaquinone |
| Colony color | Cream/white | golden | Some golden |
| Auxotrophism | Menadione, hemin, and/or thymidine | Menaquinone | Not detected |
| ATP production | Low | normal | low |

The mechanisms behind the differential effect of DGL13K and LGL13K on S. aureus resistance remain to be elucidated. The finding that DGL13K induced resistance to the stereo-isomer LGL13K but not to DGL13K itself, suggests that peptide primary structure is responsible for inducing bacterial defense mechanisms but the peptide secondary structure determines if the defense mechanisms are effective against each peptide.

## Abbreviations

AMP: antimicrobial peptide
OD: optical density
SCV: small colony variant
THB: Todd Hewitt Broth

## Acknowledgements

Dr. Juan E. Abrahante Llorens is thanked for assistance with genome sequence analysis. This work was supported by resources at the University of Minnesota Genomics Center and a UMGC Pilot Sequencing award, which are gratefully acknowledged.

